# Predicting Longitudinal Disease Severity for Individuals with Parkinson’s Disease using Functional MRI and Machine Learning Prognostic Models

**DOI:** 10.1101/2020.04.20.050971

**Authors:** Kevin P. Nguyen, Vyom Raval, Alex Treacher, Cooper Mellema, Frank Yu, Marco C. Pinho, Rathan M. Subramaniam, Richard B. Dewey, Albert Montillo

## Abstract

Parkinson’s disease is the second most common neurodegenerative disorder and is characterized by the loss of ability to control voluntary movements. Predictive biomarkers of progression in Parkinson’s Disease are urgently needed to expedite the development of neuroprotective treatments and facilitate discussions about disease prognosis between clinicians and patients. Resting-state functional magnetic resonance imaging (rs-fMRI) shows promise in predicting progression, with derived measures, including regional homogeneity (ReHo) and fractional amplitude of low frequency fluctuations (fALFF), having been previously been associated with *current* disease severity. In this work, ReHo and fALFF features from 82 Parkinson’s Disease subjects are used to train machine learning predictors of baseline clinical severity and progression at 1 year, 2 years, and 4 years follow-up as measured by the Movement Disorder Society Unified Depression Rating Scale (MDS-UPDRS) score. This is the first time that rs-fMRI and machine learning have been combined to predict *future* disease progression. The machine learning models explain up to 30.4% (*R*^2^ = 0.304) of the variance in baseline MDS-UPDRS scores, 55.8% (*R*^2^ = 0.558) of the variance in year 1 scores, and 47.1% (*R*^2^ = 0.471) of the variance in year 2 scores with high statistical significance (*p <* 0.0001). For distinguishing high- and low-progression individuals (MDS-UPDRS score above or below the median), the models achieve positive predictive values of up to 71% and negative predictive values of up to 84%. The models learn patterns of ReHo and fALFF measures that predict better and worse prognoses. Higher ReHo and fALFF in regions of the default motor network predicted lower current severity and lower future progression. The rs-fMRI features in the temporal lobe, limbic system, and motor cortex were also identified as predictors. These results present a potential neuroimaging biomarker that accurately predicts progression, which may be useful as a clinical decision-making tool and in future trials of neuroprotective treatments.

## 1. Introduction

Parkinson’s disease (PD) is the second most prevalent neurodegenerative disease, affecting 1% of individuals over the age of 60 [1]. Disease presentation and progression is highly variable among individuals, and there is no clinically accepted neurophysiological biomarker for accurately quantifying disease severity or predicting future disease progression [2, 3]. Such a biomarker would not only assist physicians in counseling patients about their disease prognosis, but also improve the design and execution of clinical trials of neuroprotective treatments. In such trials, treatment outcomes are typically quantified using clinical assessments, predominantly measuring motor symptoms, which are subjective, can be confounded by the effect of treatment, and may not truly reflect neuroprotective effects [2, 3]. An additional challenge comes from the heterogeneity of disease progression among individuals, which also confounds clinical assessments of treatment efficacy [3]. Therefore, a biomarker is needed to reliably evaluate disease progression throughout clinical trials, to provide an unbiased assessment of treatment efficacy, and to stratify subjects to expedite the enrollment of fast-progressing individuals likely to show change during the duration of the trial. A tool that can accurately predict *future* progression would empower trials to identify effective therapies, and thereby hasten the development of a treatment for Parkinson’s that slows, halts, or even reverses progression. Such a tool would also enable physicians to counsel their patients about their prognoses providing them a timeline to prepare for future needs and disabilities. Finally, an accurate predictive tool may reveal new insights into the neural correlates of disease progression.

To address this need, this work sought to develop accurate neuroimaging biomarkers of disease severity and progression taking a machine learning-based approach to the assessment of resting-state functional magnetic resonance imaging (rs-fMRI). Previous studies have identified associations between measures derived from rs-fMRI and disease severity. Regional homogeneity (ReHo) quantifies the similarity of each voxel’s activity with that of its neighbors [4]. ReHo in the cerebellum and lingual gyrus has been shown to be positively correlated with disease severity, while ReHo in the putamen is negatively correlated with Parkinson’s disease severity [5, 6]. Amplitude of low frequency fluctuation (ALFF) and its normalized form, fractional ALFF (fALFF), are measures of the power of the low frequency resting-state signals [7, 8]. ALFF in the left inferior parietal lobe, bilateral precuneus, and the right inferior frontal gyrus pars opercularis and fALFF in the right cerebellum have been positively correlated with disease severity [9, 10, 11]. Conversely, ALFF in the right prefrontal cortex, right middle occipital gyrus, and bilateral lingual gyri have all been negatively correlated with PD severity [9]. These associations are shown in **Figure 1**.

**Figure 1:**
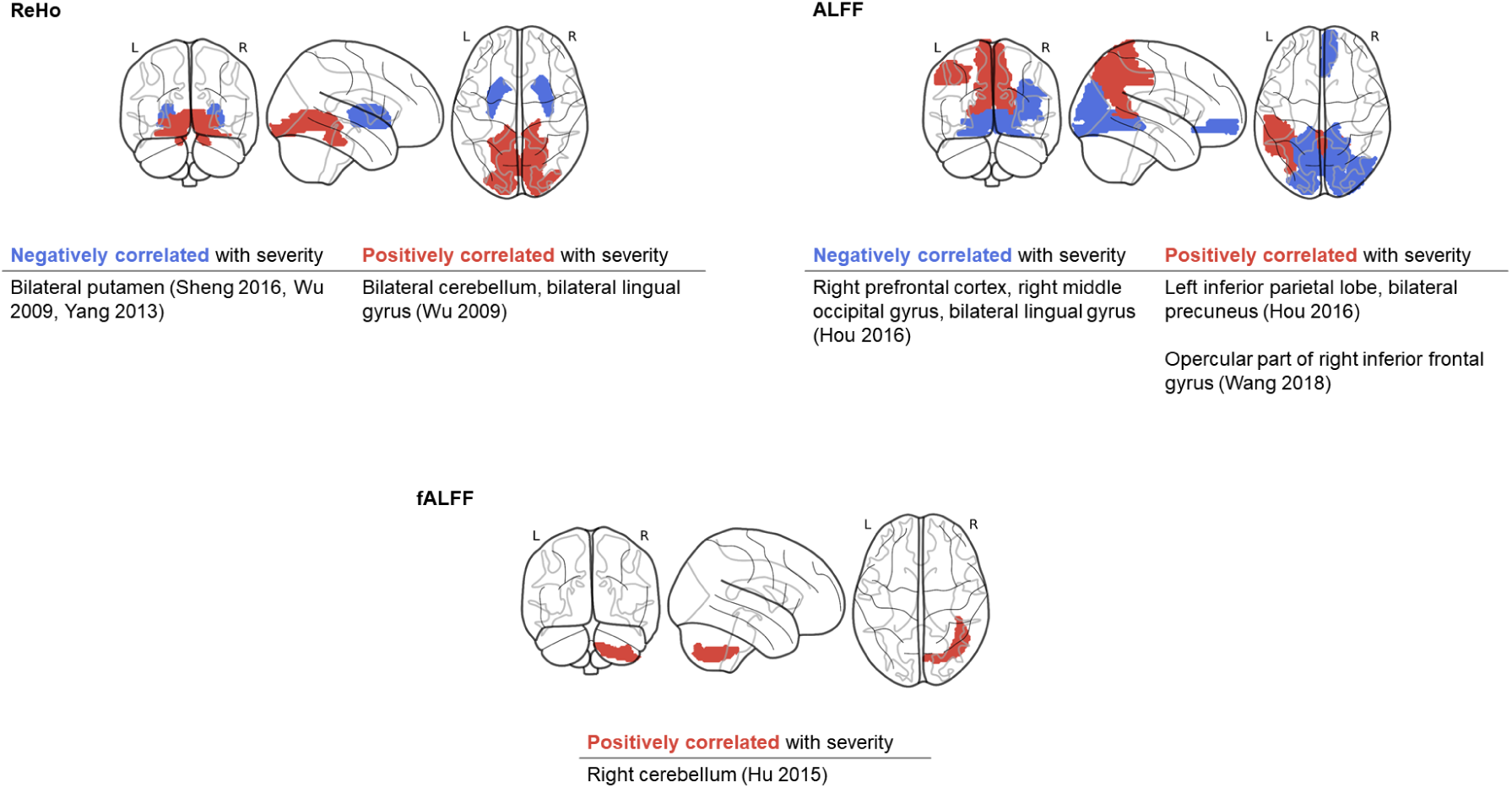
Resting-state functional MRI measurements previously found to be associated with current disease severity, visualized qualitatively. Red regions indicate positive correlations with severity, while blue regions indicate negative correlations.

Building upon these previous findings, the contributions of this work are as follows: *First*, this work aims to develop predictors of *future* progression, which may reveal findings more relevant to the pathophysiology of PD than imaging biomarkers that are correlated solely with baseline severity. *Second*, this work adopts and extensively optimizes a multivariate, machine learning approach. With higher statistical capacity, machine learning models are more likely to find complex associations between neuroimaging features and progression than the voxel-wise or univariate approaches which have been the predominant approach in the literature [5, 6, 10, 11]. Through cross-validation, the accuracy of the predictive models is rigorously evaluated on held-out data, providing more confidence that the associations found will generalize to subjects beyond the current dataset. One previous study has combined ALFF and machine learning using a relevance vector machine to predict current Movement Disorder Society Unified Parkinson’s Disease Rating Scale (MDS-UPDRS) Part III score, a measure of motor severity [9]. In comparison, this work aims to also investigate ReHo, evaluate a wider range of machine learning models, and predict both current and future MDS-UPDRS Total score. *Finally*, in this work the predictive power of neural correlates from ReHo and fALFF are compared, as well as the specific brain regions that were found to be most important for prediction. These predictive regional imaging features comprise a candidate composite neuroimaging biomarker of PD progression.

## 2. Methods

### 2.1. Dataset

Data used in the preparation of this article were obtained from the Parkinson’s Progression Markers Initiative (PPMI) database (www.ppmi-info.org/data). The outcome of interest in this analysis is the MDS-UPDRS total score, encompassing both motor and non-motor symptomatology. From this database, 82 PD subjects with rs-fMRI and outcome scores at the same visit were acquired for analysis. This initial imaging visit is considered the *baseline* visit for this analysis. Of these 82 subjects, 53 subjects also had outcome scores available at year 1 after imaging, 45 subjects had scores available at year 2 after imaging, and 33 had scores at year 4 after imaging. MDS-UPDRS scores included the Part III Motor Examination conducted in the on-medication state. Off-medication scores were not used because 1) they were not available for over half of the subjects and 2) the on-medication scores are more reflective of the underlying dopaminergic *and* non-dopaminergic neuropathology despite best possible correction with dopaminergic medication. On-medication measurements of severity are therefore more relevant to neuroprotective treatment development than off-medication measurements, which are dominated by symptoms of dopaminergic neuron loss. The distributions of MDS-UPDRS at baseline, year 1, year 2, and year 4 are illustrated in **Figure 2**. The median MDS-UPDRS score at each timepoint was used to separate subjects into high- and low-progression groups. The median score was 32 at baseline, 35 at year 1, 37 at year 2, and 36 at year 4. This indicates a trend of increasing severity with time, except for year 4 which has a limited number of subjects and showed wider variance in MDS-UPDRS score.

**Figure 2:**
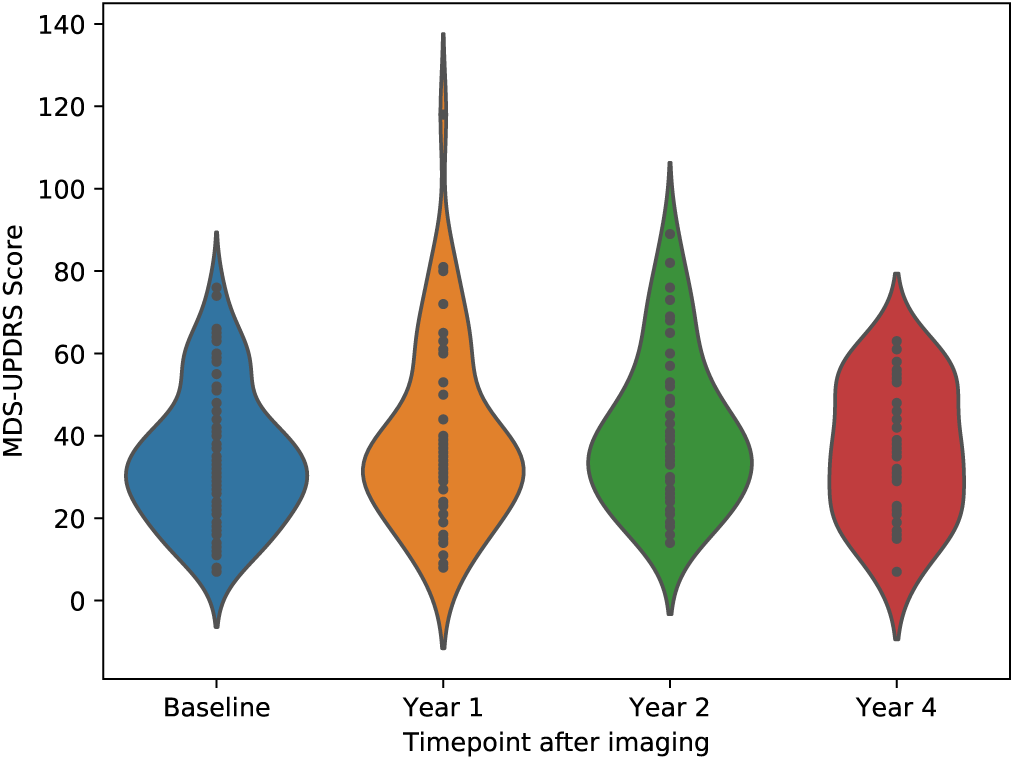
Distribution of MDS-UPDRS scores at each timepoint.

fMRI was acquired at resting-state on 3T Siemens scanners at 8 study sites, using a 2D echo planar imaging sequence with TR 2400 ms, TE 25 ms, flip angle 80°, and 3.3 mm isotropic voxels. Final image dimensions were 68 × 66 × 33 voxels with 210 volumes. Total acquisition time was 504 seconds.

In addition to fMRI, clinical and demographic features were included as covariates during training of the predictive models. Clinical features included disease duration, symptom duration, dominant symptom side, Geriatric Depression Scale (GDS), Montreal Cognitive Assessment (MoCA), and presence of tremor, rigidity, bradykinesia, or postural instability (encoded as binary variables) at baseline. Baseline MDS-UPDRS score was also included as a confounding variable when training models to predict longitudinal outcomes. Demographic features included age, sex, ethnicity, race, handedness, and years of education. These clinical and demographic characteristics are summarized in **Table 1**.

**Table 1:**
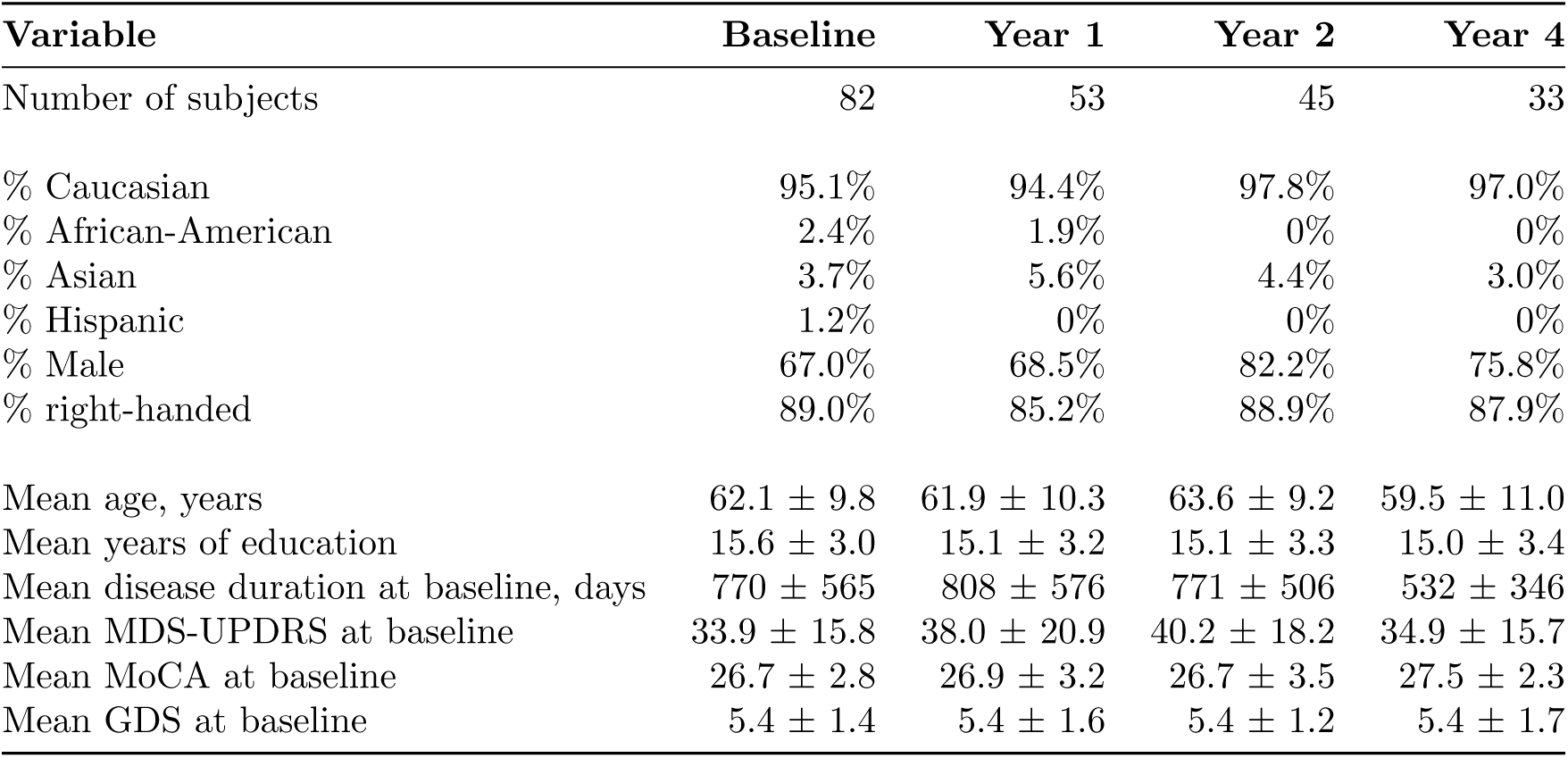
Demographics and baseline disease characteristics of subjects included in this analysis.

| Variable | Baseline | Year 1 | Year 2 | Year 4 |
| --- | --- | --- | --- | --- |
| Number of subjects | 82 | 53 | 45 | 33 |
| % Caucasian | 95.1% | 94.4% | 97.8% | 97.0% |
| % African-American | 2.4% | 1.9% | 0% | 0% |
| % Asian | 3.7% | 5.6% | 4.4% | 3.0% |
| % Hispanic | 1.2% | 0% | 0% | 0% |
| % Male | 67.0% | 68.5% | 82.2% | 75.8% |
| % right-handed | 89.0% | 85.2% | 88.9% | 87.9% |
| Mean age, years | 62.1 ± 9.8 | 61.9 ± 10.3 | 63.6 ± 9.2 | 59.5 ± 11.0 |
| Mean years of education | 15.6 ± 3.0 | 15.1 ± 3.2 | 15.1 ± 3.3 | 15.0 ± 3.4 |
| Mean disease duration at baseline, days | 770 ± 565 | 808 ± 576 | 771 ± 506 | 532 ± 346 |
| Mean MDS-UPDRS at baseline | 33.9 ± 15.8 | 38.0 ± 20.9 | 40.2 ± 18.2 | 34.9 ± 15.7 |
| Mean MoCA at baseline | 26.7 ± 2.8 | 26.9 ± 3.2 | 26.7 ± 3.5 | 27.5 ± 2.3 |
| Mean GDS at baseline | 5.4 ± 1.4 | 5.4 ± 1.6 | 5.4 ± 1.2 | 5.4 ± 1.7 |

### 2.2. Preprocessing

fMRI images were first realigned to the mean volume with affine transformations to correct for inter-volume head motion, using the MCFLIRT tool in FSL [12]. Brain masking was performed using AFNI 3dAutomask [13]. The images were next nonlinearly coregistered directly to a common EPI template in MNI space, because direct EPI spatial normalization has been shown to correct for more nonlinear magnetic susceptibility artifacts than T1-based normalization [14, 15]. This nonlinear normalization step was performed using the Symmetric Normalization algorithm in ANTS [16].

Because individuals with PD may exhibit substantial head motion during imaging, it is crucial to correct for head motion artifacts in fMRI data. Left uncorrected, these artifacts may introduce spurious and confounding signals into subsequent analysis. Motion-related signals in the fMRI data were identified using ICA-AROMA [17]. Recent work suggests that all fMRI nuisance regression should be performed simultaneously to avoid reintroduction of noise [18]. Accordingly, such an approach has been developed and applied here in which the motion-related regressors computed by ICA-AROMA are concatenated with the affine head motion parameters computed during inter-volume realignment and the mean white matter and cerebrospinal fluid signals, and these nuisance signals were regressed out of the fMRI data in one step [19].

Regional homogeneity (ReHo), amplitude of low frequency and fractional amplitude of low frequency fluctuations (fALFF) were computed from the cleaned fMRI using C-PAC [20]. ReHo was computed using Kendall’s coefficient of concordance between each voxel and its 27-voxel neighborhood. Low frequency power was measured by applying linear detrending and bandpass filtering at 0.01–0.1 Hz to each voxel’s signal, then computing the standard deviation of the signal. This was divided by the standard deviation of the unfiltered signal to obtain fALFF. To normalize the values, the Z-scores for ReHo and fALFF were calculated per subject.

### 2.3. Model Training and Evaluation

To extract regional features from the ReHo and fALFF maps, three different brain parcellations of varying granularity were applied. Parcellations with a higher number of regions-of-interest (ROIs) yield more spatially precise but potentially more noisy features, and this trade-off was investigated by testing two different parcellations. These included the 100-ROI Schaefer functional brain parcellation, modified with an additional 35 striatal and cerebellar ROIs [21] and the 197-ROI Bootstrap Analysis of Stable Clusters (BASC197) atlas [22]. These parcellations were used to compute the mean regional ReHo or fALFF values for each subject.

To test the ability of ReHo and fALFF features to predict current and future disease severity, four prediction targets were examined: MDS-UPDRS total score at baseline, year 1, year 2, and year 4. Separate machine learning models were trained and optimized to predict each target from either ReHo or fALFF features, with clinical and demographic features added as covariates. Four machine learning models of varying statistical complexity were tested for each target-feature combination: ElasticNet regression, Support Vector Machine (SVM) with a linear kernel, Random Forest with a decision tree kernel, and Gradient Boosting with a decision tree kernel. To determine the best-performing parcellation and model combination for each target, a nested cross-validation approach was applied. A leave-one-out cross-validation (LOOCV) approach was adopted for the outer cross-validation loop, with one subject held out of the data and the remaining subjects used for subsequent model and parcellation selection. Hyperparameters for each of the 4 models were optimized through a random search of 300 hyperparameter and parcellation configurations (100 hyperparameter configurations per parcellation) with an inner 10-fold cross-validation loop. The best performing model-hyperparameter-parcellation combination was selected based on root mean squared error (RMSE), the model was retrained using all training and validation data, and predictive performance was evaluated on the held-out subject. This process was repeated until all subjects had been held out once. The final predictive performance on the held-out subjects is reported in the results. Each model’s predictions were also thresholded *post-hoc* to evaluate the model’s ability to classify high-versus low-progression subjects.

### 2.4. Feature Importance Analysis

The high performing models for each prediction target were examined to determine which features were learned as important predictors of progression. For the ElasticNet and linear-kernel SVM models, feature importance was determined from the coefficients of the trained models, where coefficients of higher magnitude indicate more important features. The sign of the coefficient then indicates whether the feature is positively or negatively associated with the prediction target. For the Random Forest and Gradient Boosting models, feature importance was determined using Gini importance (mean decrease in impurity) [23]. After computing feature importance at each iteration of the LOOCV loop, the median importance was obtained for each feature. As this is an unsigned value, univariate linear correlation was used to determine the direction of association between each feature and the prediction target.

To aid with interpretation and comparison between different brain parcellations, anatomical labels were assigned to the imaging features using the Automated Anatomical Labeling (AAL) atlas [24]. Specifically, the centroid of each feature’s ROI was computed, and the feature was assigned the nearest anatomical label in the AAL atlas.

## 3. Results

### 3.1. Predictive Performance

Predictive performance results for each of the four MDS-UPDRS targets are summarized in **Table 2**. ReHo features explained 38.0%, 46.8%, 51.2%, and 25.2% of the variance in baseline, year 1, year 2, and year 4 MDS-UPDRS score, respectively. fALFF features explained 23.3%, 56.6%, 40.2%, and 7.5% of the variance in baseline, year 1, year 2, and year 4 MDS-UPDRS score, respectively. **Figure 3** plots the model prediction versus ground truth MDS-UPDRS scores for the two timepoints, years 1 and 2, at which predictive performance was highest. For classifying high-versus low-progression subjects, ReHo and fALFF features achieved similar accuracy, with positive predictive value (PPV) ranging from 60.9% to 71.4%.

**Table 2:** Predictive performance achieved for each MDS-UPDRS timepoint and each imaging feature type, computed through leave-one-out cross-validation. Metrics for classifying high-progression subjects (AUC, PPV, NPV) were computed by thresholding predicted and ground truth MDS-UPDRS scores at the median, dichotomizing subjects into high- and low-progression groups. Metrics: *R*^2^, coefficient of determination; RMSE: root mean squared error, AUC: area under the receiver operating characteristic curve, PPV: positive predictive value, NPV: negative predictive value.

| MDS-UPDRS<br>Prediction target | Feature | Best performing model | Best performing<br>parcellation | $R^2$ ($p$-value) | RMSE | AUC | PPV | NPV |
| --- | --- | --- | --- | --- | --- | --- | --- | --- |
| Baseline | ReHo | Gradient Boosting | Schaefer | 0.304 ($p < 0.0001$) | 13.415 | 0.683 | 66.7% | 70.0% |
| | fALFF | Gradient Boosting | Schaefer | 0.242 ($p < 0.0001$) | 14.006 | 0.708 | 69.0% | 72.5% |
| Year 1 | ReHo | ElasticNet | Schaefer | 0.453 ($p < 0.0001$) | 15.861 | 0.757 | 71.4% | 80.0% |
| | fALFF | ElasticNet | Schaefer | 0.558 ($p < 0.0001$) | 14.256 | 0.757 | 71.4% | 80.0% |
| Year 2 | ReHo | ElasticNet | Schaefer | 0.471 ($p < 0.0001$) | 13.322 | 0.717 | 65.4% | 78.9% |
| | fALFF | ElasticNet | Schaefer | 0.463 ($p < 0.0001$) | 13.426 | 0.717 | 69.2% | 84.2% |
| Year 4 | ReHo | SVM | BASC197 | 0.255 ($p = 0.003$) | 14.015 | 0.663 | 60.9% | 68.2% |
| | fALFF | SVM | BASC197 | 0.152 ($p = 0.025$) | 14.957 | 0.667 | 70.0% | 72.0% |

**Figure 3:**
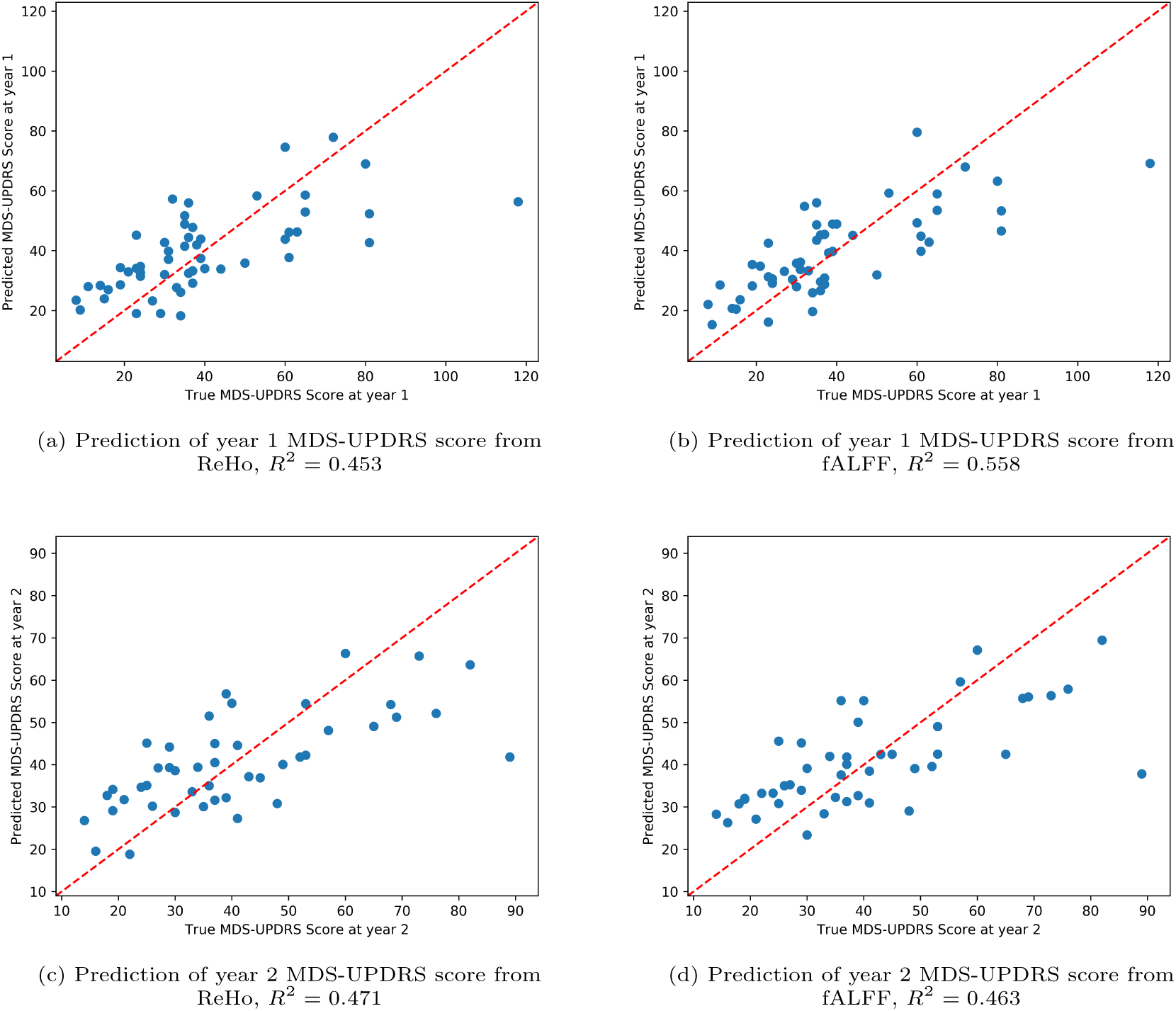
Predicted versus ground truth scores for (a,c) ReHo and (b,d) fALFF for top models.

### 3.2. Predictive ReHo Features

At baseline (**Figure 4a**), ReHo in the left orbitofrontal cortex, right posterior and middle cingulate gyri, and bilateral precuneus were important regions predictive of lower MDS-UPDRS score. Greater disease duration (time since diagnosis) and symptom duration were predictive of higher score. At year 1 after imaging (**Figure 4b**), while baseline MDS-UPDRS was the strongest predictor of year 1 MDS-UPDRS score, three imaging features were also found to be important. ReHo in the right inferior frontal gyrus pars triangularis and left amygdala were predictive of lower score while ReHo in the right angular gyrus was predictive of higher score. At year 2 (**Figure 4c**), baseline MDS-UPDRS was a strong predictor of higher score as were imaging measures. In particular, ReHo in several regions in the right inferior temporal lobe as well as the right putamen and posterior cingulate gyrus were predictive of lower score. At year 4 (**Figure 4d**), ReHo in the left orbitofrontal cortex, right inferior frontal gyrus pars triangularis, left Rolandic operculum, right middle temporal gyrus, and left thalamus were predictive of lower score. ReHo in the middle frontal gyrus and inferior temporal gyrus were important predictors of higher score at year 4. Baseline MDS-UPDRS was not found to be an important predictor of year 4 score.

**Figure 4:**
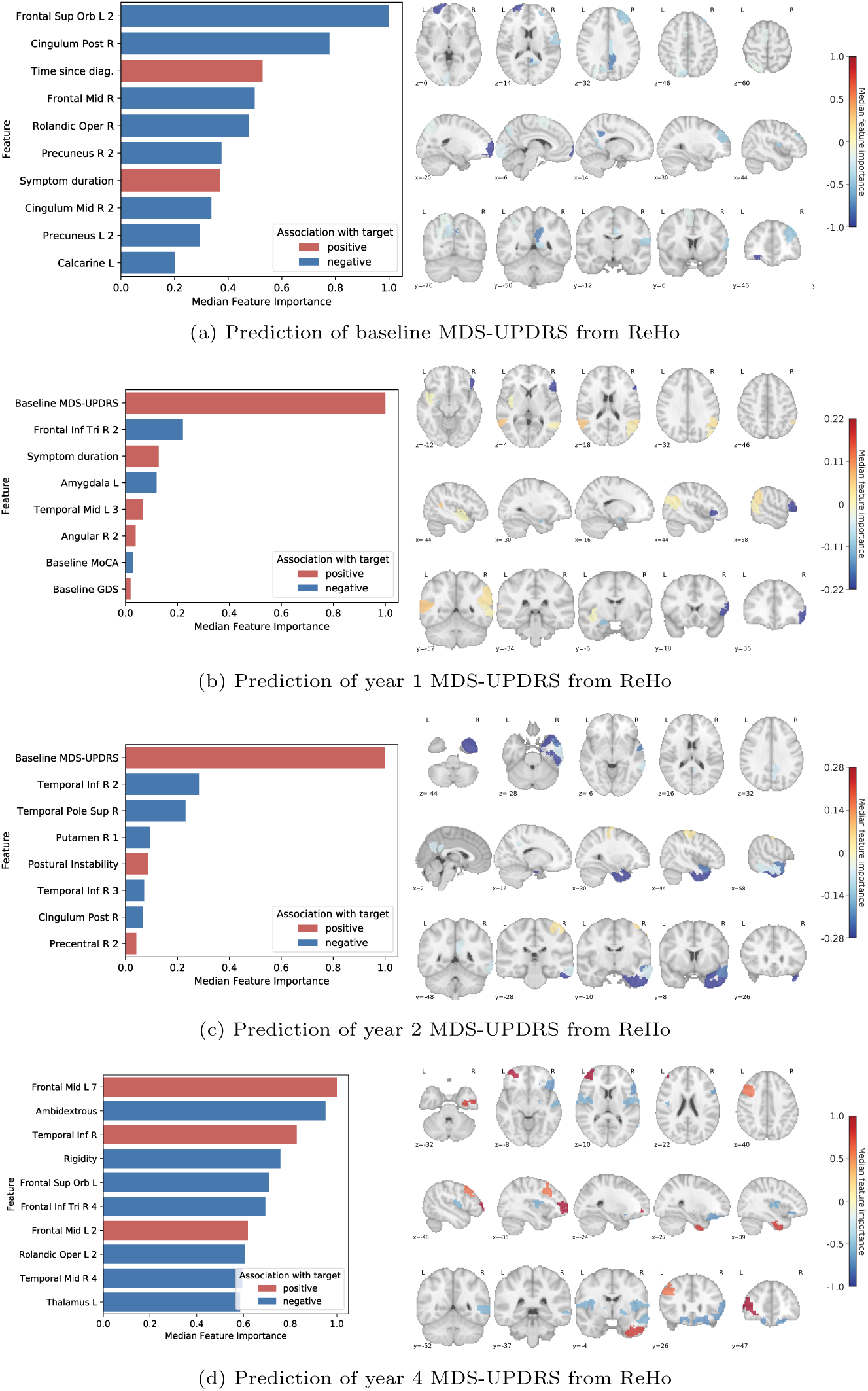
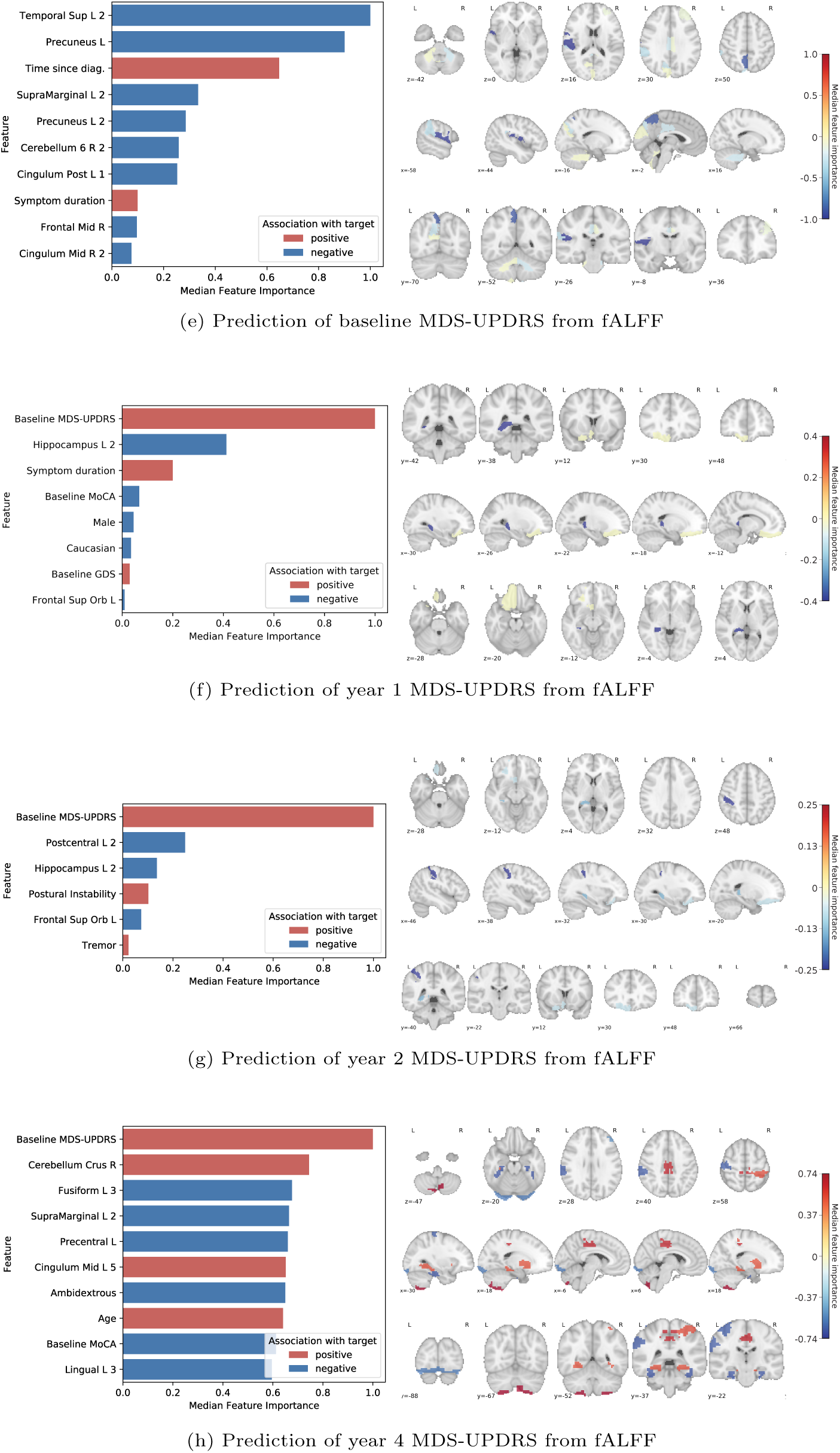
Important features learned by each model to predict MDS-UPDRS score. The median feature importance among the LOOCV iterations is shown. *Left*: the most important features are illustrated, sorted by absolute importance. For brevity, features with zero or comparatively low importance are not shown. Red bars indicate a positive association between the feature and MDS-UPDRS score, and blue bars indicate a negative association. Importance values are normalized to the range of 0–1. *Right*: the imaging features are visualized in brain space, overlaid on a standard MNI template.

### 3.3. Predictive fALFF Features

At baseline (**Figure 4e**), fALFF in the left superior temporal gyrus, left precuneus, left supramarginal gyrus, right cerebellar lobe 6, and bilateral cingulate gyrus were predictive of lower MDS-UPDRS score. Greater disease duration and symptom duration were again predictive of higher score. At year 1 (**Figure 4f**), baseline MDS-UPDRS was the strongest predictor of MDS-UPDRS score, while fALFF in the left hippocampus was predictive of lower score. At year 2 (**Figure 4g**), fALFF in the left postcentral gyrus, left hippocampus, and left orbitofrontal cortex were predictive of lower score. Baseline MDS-UPDRS, presence of a tremor, and postural instability predicted higher year 2 score. Finally, at year 4 (**Figure 4h**), fALFF in the left fusiform gyrus, left supramarginal gyrus, left precentral gyrus, and left lingual gyrus were predictive of lower score. fALFF in the right cerebellum and left middle cingulate gyrus were predictive of higher score.

## 4. Discussion

### 4.1. Performance of Predictive Models

The machine learning models trained to predict baseline and longitudinal MDS-UPDRS scores from ReHo and fALFF features performed better than previously proposed models. To date, Hou et al. [9] were the first to use machine learning to predict PD severity from rs-fMRI measurements. They achieved *R*^2^ = 0.123 for predicting UPDRS Part III score using ALFF features and a relevance vector regressor model. In comparison, the models in this work achieved *R*^2^ = 0.304, *p <* 0.0001 using ReHo and *R*^2^ = 0.242, *p <* 0.0001 using fALFF for predicting MDS-UPDRS total score. While the Part III score measures motor symptoms, the total score includes both motor and non-motor symptoms and is a more comprehensive assessment of PD severity. The models developed in this work also explained a substantial percentage of the variance in MDS-UPDRS scores at years 1 and 2 where *R*^2^ ranged from 0.453 to 0.558, *p <* 0.0001, and such future progression has not been addressed previously using these fMRI measures. The addition of baseline MDS-UPDRS score as a feature at these timepoints was a contributor to the high predictive accuracy, though this feature became less important at further timepoints. Performance was lower though still substantial for year 4 (*R*^2^ = 0.255, *p* = 0.003 for ReHo features and *R*^2^ = 0.152, *p* = 0.025 for fALFF features). This could be attributed to either the difficulty of predicting long-term progression or the limited number of available subjects at this timepoint (*n* = 33). Examination of predicted versus true MDS-UPDRS scores (**Figure 3**) showed similar accuracy throughout the range of low to moderate severity subjects. The greatest prediction errors occurred for outlier subjects with very high MDS-UPDRS scores.

When dichotomizing the subjects into high- and low-progression using the median MDS-UPDRS score as the threshold, the year 1 and year 2 models achieved high NPVs of 80.0% to 85.7% in identifying high-progression subjects. PPVs were comparatively lower, ranging from 65.4% to 71.4%. The baseline models were relatively less capable of this classification task, and PPV and NPV were lower for the year 4 models, indicative of the difficulty of predicting very long range disease trajectory.

In additional experiments not elaborated here, models were trained on combined ReHo and fALFF features. There was no significant increase in performance, suggesting that at least some of the features from the two measurements are collinear and measure similar underlying signals. Further analysis to identify the most informative and synergistic features from ReHo and fALFF is warranted.

### 4.2. Learned Predictive Features

Across the timepoints, ReHo in many regions associated with the default mode network (DMN) were predictive of lower severity or progression, including the left orbitofrontal cortex, bilateral prefrontal cortex, right cingulate gyrus, and bilateral precuneus. The DMN is a well characterized resting-state network with known roles in cognitive processing and executive function [25], and disruptions in this network have been identified in PD [26, 27]. These findings suggest that higher neural homogeneity in the DMN predicts lower disease progression. ReHo in the right inferior frontal gyrus pars triangularis was a highly important predictor of lower progression at years 1 and 4. This region contains Brodmann Area 45 (Broca’s area), which is known to be involved in fine hand movement [28], and hand tremors are a hallmark of PD. While not extensively implicated in PD, one study has correlated cortical atrophy in Broca’s area with disease duration, suggesting a role of this region in progression [29]. Higher neural homogeneity in the pars triangularis could potentially be an important indicator of lower progression, particularly in hand tremor symptoms. Another recurring important region was the right temporal pole and nearby areas of the right middle and inferior temporal gyri, where ReHo was predictive of year 2 and 4 progression. Gray matter atrophy in the temporal pole, which exhibits functional connections to the nearby striatum and orbitofrontal cortex, has been identified in PD [30], correlated with disease duration [29], and connected to cognitive impairment [31]. Importantly, the anterior temporal areas are the site of initial cortical involvement in Braak’s model of progression in PD [32].

At baseline, fALFF in several regions of the DMN was predictive of lower severity, including the left precuneus and right posterior and middle cingulate gyri. The left orbitofrontal cortex, another component of the DMN, was a recurring predictive region of progression at years 1 and 2. Combined with similar findings in the ReHo features, this strongly implicates higher DMN activity as an indicator of lower progression. Higher fALFF in the left superior temporal and supramarginal gyri predicted lower baseline severity, specifically in ROIs located close to the insula. Pathological involvement of the insula, a component of the limbic system, has been suggested to be a contributor to cognitive decline [33, 34]. Higher fALFF in the left hippocampus, another limbic region, predicted lower short-term progression at years 1 and 2. Indeed, limbic system involvement is characteristic of progressive disease in the Braak’s model [35]. Finally, fALFF in motor regions such as the right cerebellum and left postcentral gyrus were predictors of both baseline severity and longer-term (years 2 and 4) progression.

### 4.3. Limitations

A limitation to this work is the sample size of the PPMI dataset used to train the predictive models, particularly for longer range predictions. The initial cohort contained 82 subjects at baseline. However, subject dropout led to fewer available subjects at each successive timepoint, with 33 subjects remaining at year 4. Though we have used more subjects than previous studies which have successfully detected associations between rs-fMRI and PD severity/progression with 39, 22, and even 17 subjects [6, 5, 36], we expect a larger dataset would likely increase the predictive power of the models. Furthermore, the group of 33 subjects at year 4 had earlier-stage disease (mean disease duration 532 ± 346 versus 770 ± 565 days, *p* = 0.03) compared to the original 82 subjects. The mean age was also lower (59.5 ± 11.0 versus 62.1 ± 9.8 years), but the difference was not significant (*p* = 0.22). An additional limitation of the dataset is the bias towards Caucasians (95.1%) and college-educated individuals (mean years of education 15.6 ± 3.0).

## 5. Conclusion

Using a multivariate machine learning approach, predictors of progression were developed that explained a large percentage of the variance in MDS-UPDRS score in PD subjects over multiple timepoints. ReHo and fALFF were demonstrated to have prognostic value, containing a predictive signal capable of *quantitatively* forecasting progression for up to 4 years. The machine learning models were examined to identify important imaging and clinical features, which together form a composite biomarker of disease progression. Higher ReHo and fALFF in regions of the default motor network were found to predict both lower baseline severity and lower progression. ReHo in the inferior frontal gyrus pars triangularis (Broca’s area) was a predictor of short-term progression, while temporal lobe ReHo was implicated in longer-term progression. Additionally, fALFF in limbic and motor regions were identified as predictive of progression. Baseline MDS-UPDRS score was a strong predictor of 1- and 2-year progression but less predictive of 4-year progression.

These results can help to fulfill the urgent need for accurate and predictive biomarkers to facilitate the development of neuroprotective treatments. This work provides strong evidence of the value of rs-fMRI in prognosticating PD, and continued efforts to develop imaging-based tools will not only expedite treatment development, but also improve understanding of PD neurophysiology and patient care.

## Acknowledgements

PPMI-–a public-private partnership-–is funded by the Michael J. Fox Foundation for Parkinson’s Research and funding partners, including Abbvie, Allergan, Avid Radiopharmaceuticals, Biogen, Biolegend, Bristol-Myers Squibb, Celgene, Denali, GE Healthcare, Genentech, GlaxoSmithKline, Lilly, Lundbeck, Merck, Meso Scale Discovery, Pfizer, Piramal, Prevail Therapeutics, Roche, Sanofi Genzyme, Servier, Takeda, Teva, UCB, Verily, and Voyager Therapeutics.

## Notes

### Competing Interest Statement

The authors have declared no competing interest.

## References

[1] T. Pringsheim, N. Jette, A. Frolkis, T. D. L. Steeves, The prevalence of Parkinson’s disease: a systematic review and meta-analysis, Movement disorders: official journal of the Movement Disorder Society 29 (13) (2014) 1583–1590. doi:10.1002/mds.25945.

[2] W. G. Meissner, M. Frasier, T. Gasser, C. G. Goetz, A. Lozano, P. Piccini, J. A. Obeso, O. Rascol, A. Schapira, V. Voon, D. M. Weiner, F. Tison, E. Bezard, Priorities in Parkinson’s disease research, Nature reviews. Drug discovery 10 (5) (2011) 377–393. doi:10.1038/nrd3430.

[3] K. Gwinn, K. K. David, C. Swanson-Fischer, R. Albin, C. S. Hillaire-Clarke, B.-A. Sieber, C. Lungu, F. D. Bowman, R. N. Alcalay, D. Babcock, T. M. Dawson, R. B. Dewey, T. Foroud, D. German, X. Huang, V. Petyuk, J. A. Potashkin, R. Saunders-Pullman, M. Sutherland, D. R. Walt, A. B. West, J. Zhang, A. Chen-Plotkin, C. R. Scherzer, D. E. Vaillancourt, L. S. Rosenthal, Parkinson’s disease biomarkers: perspective from the NINDS Parkinson’s Disease Biomarkers Program, Biomarkers in medicine 11 (6) (2017) 451–473. doi:10.2217/bmm-2016-0370.

[4] Y. Zang, T. Jiang, Y. Lu, Y. He, L. Tian, Regional homogeneity approach to fMRI data analysis, NeuroImage 22 (1) (2004) 394–400. doi:10.1016/j.neuroimage.2003.12.030.

[5] T. Wu, X. Long, Y. Zang, L. Wang, M. Hallett, K. Li, P. Chan, Regional homogeneity changes in patients with parkinson’s disease, Human brain mapping 30 (5) (2009) 1502–1510. doi:10.1002/hbm.20622.

[6] K. Sheng, W. Fang, Y. Zhu, G. Shuai, D. Zou, M. Su, Y. Han, O. Cheng, Different alterations of cerebral regional homogeneity in early-onset and late-onset Parkinson’s disease, Frontiers in Aging Neuroscience 8 (JUN) (2016). doi:10.3389/fnagi.2016.00165.

[7] Y.-F. Zang, Y. He, C.-Z. Zhu, Q.-J. Cao, M.-Q. Sui, M. Liang, L.-X. Tian, T.-Z. Jiang, Y.-F. Wang, Altered baseline brain activity in children with ADHD revealed by resting-state functional MRI, Brain & development 29 (2) (2007) 83–91. doi:10.1016/j.braindev.2006.07.002.

[8] Q.-H. Zou, C.-Z. Zhu, Y. Yang, X.-N. Zuo, X.-Y. Long, Q.-J. Cao, Y.-F. Wang, Y.-F. Zang, An improved approach to detection of amplitude of low-frequency fluctuation (ALFF) for resting-state fMRI: fractional ALFF, Journal of neuroscience methods 172 (1) (2008) 137–141. doi:10.1016/j.jneumeth.2008.04.012.

[9] Y. Hou, C. Luo, J. Yang, R. Ou, W. Song, Q. Wei, B. Cao, B. Zhao, Y. Wu, H.-F. Shang, Q. Gong, Prediction of individual clinical scores in patients with Parkinson’s disease using resting-state functional magnetic resonance imaging, Journal of the Neurological Sciences 366 (2016) 27–32. doi:10.1016/j.jns.2016.04.030.

[10] Z. Wang, X. Jia, H. Chen, T. Feng, H. Wang, Abnormal Spontaneous Brain Activity in Early Parkinson’s Disease With Mild Cognitive Impairment: A Resting-State fMRI Study, Frontiers in physiology 9 (2018) 1093. doi:10.3389/fphys.2018.01093.

[11] X.-F. Hu, J.-Q. Zhang, X.-M. Jiang, C.-Y. Zhou, L.-Q. Wei, X.-T. Yin, J. Li, Y.-L. Zhang, J. Wang, Amplitude of low-frequency oscillations in Parkinson’s disease: A 2-year longitudinal resting-state functional magnetic resonance imaging study, Chinese Medical Journal 128 (5) (2015) 593–601. doi:10.4103/0366-6999.151652.

[12] M. Jenkinson, P. Bannister, M. Brady, S. Smith, Improved Optimization for the Robust and Accurate Linear Registration and Motion Correction of Brain Images, NeuroImage 17 (2) (2002) 825–841. doi:10.1016/S1053-8119(02)91132-8.

[13] R. W. Cox, AFNI: Software for Analysis and Visualization of Functional Magnetic Resonance Neuroimages, Computers and Biomedical Research 29 (3) (1996) 162–173. doi:10.1006/cbmr.1996.0014.

[14] V. D. Calhoun, T. D. Wager, A. Krishnan, K. S. Rosch, K. E. Seymour, M. B. Nebel, S. H. Mostofsky, P. Nyalakanai, K. Kiehl, The impact of T1 versus EPI spatial normalization templates for fMRI data analyses, Human brain mapping 38 (11) (2017) 5331–5342. doi:10.1002/hbm.23737.

[15] E. Dohmatob, G. Varoquaux, B. Thirion, Inter-subject Registration of Functional Images: Do We Need Anatomical Images?, Frontiers in Neuroscience 12 (2018). doi:10.3389/fnins.2018.00064.

[16] B. B. Avants, N. J. Tustison, G. Song, P. A. Cook, A. Klein, J. C. Gee, A Reproducible Evaluation of ANTs Similarity Metric Performance in Brain Image Registration, NeuroImage 54 (3) (2010) 2033–2044. doi:10.1016/j.neuroimage.2010.09.025.

[17] R. H. R. Pruim, M. Mennes, D. van Rooij, A. Llera, J. K. Buitelaar, C. F. Beckmann, ICA-AROMA: A robust ICA-based strategy for removing motion artifacts from fMRI data, NeuroImage 112 (2015) 267–277. doi:10.1016/j.neuroimage.2015.02.064.

[18] M. A. Lindquist, S. Geuter, T. D. Wager, B. S. Caffo, Modular preprocessing pipelines can reintroduce artifacts into fMRI data, Human brain mapping 40 (8) (2019) 2358–2376. doi:10.1002/hbm.24528.

[19] V. Raval, K. P. Nguyen, A. Montillo, Improved Motion Correction for Functional MRI using an Omnibus Regression Model, in: IEEE International Symposium on Biomedical Imaging (ISBI), IEEE, 2020. URL Https://arxiv.org/pdf/1911.10229.pdf

[20] Cameron Craddock, Sharad Sikka, Brian Cheung, Ranjeet Khanuja, Satrajit Ghosh, Chaogan Yan, Qingyang Li, Daniel Lurie, Joshua Vogelstein, Randal Burns, Stanley Colcombe, Maarten Mennes, Clare Kelly, M. Di Adriana, Francisco Castellanos, Michael Milham, Towards Automated Analysis of Connectomes: The Configurable Pipeline for the Analysis of Connectomes (C-PAC), Frontiers in Neuroinformatics 7 (2013). doi:10.3389/conf.fninf.2013.09.00042.

[21] A. Schaefer, R. Kong, E. M. Gordon, T. O. Laumann, X.-N. Zuo, A. Holmes, S. B. Eickhoff, B. T. T. Yeo, Local-Global Parcellation of the Human Cerebral Cortex From Intrinsic Functional Connectivity MRI, 2017. doi:10.1101/135632.

[22] P. Bellec, P. Rosa-Neto, O. C. Lyttelton, H. Benali, A. C. Evans, Multi-level bootstrap analysis of stable clusters in resting-state fMRI, NeuroImage 51 (3) (2010) 1126–1139. doi:10.1016/j.neuroimage.2010.02.082.

[23] L. Breiman, Random forests, Machine Learning 45 (1) (2001) 5–32. doi:10.1023/A:1010933404324.

[24] N. Tzourio-Mazoyer, B. Landeau, D. Papathanassiou, F. Crivello, O. Etard, N. Delcroix, B. Mazoyer, M. Joliot, Automated anatomical labeling of activations in SPM using a macroscopic anatomical parcellation of the MNI MRI single-subject brain, NeuroImage 15 (1) (2002) 273–289. doi:10.1006/nimg.2001.0978.

[25] M. D. Greicius, B. Krasnow, A. L. Reiss, V. Menon, Functional connectivity in the resting brain: a network analysis of the default mode hypothesis, Proceedings of the National Academy of Sciences of the United States of America 100 (1) (2003) 253–258. doi:10.1073/pnas.0135058100.

[26] T. van Eimeren, O. Monchi, B. Ballanger, A. P. Strafella, Dysfunction of the default mode network in parkinson disease: a functional magnetic resonance imaging study, Archives of neurology 66 (7) (2009) 877–883. doi:10.1001/archneurol.2009.97.

[27] A. Tessitore, F. Esposito, C. Vitale, G. Santangelo, M. Amboni, A. Russo, D. Corbo, G. Cirillo, P. Barone, G. Tedeschi, Default-mode network connectivity in cognitively unimpaired patients with parkinson disease, Neurology 79 (23) (2012) 2226–2232. doi:10.1212/WNL.0b013e31827689d6.

[28] F. Binkofski, G. Buccino, Motor functions of the Broca’s region, Brain and Language 89 (2) (2004) 362–369. doi:10.1016/S0093-934X(03)00358-4.

[29] T. Jubault, J.-F. Gagnon, S. Karama, A. Ptito, A.-L. Lafontaine, A. C. Evans, O. Monchi, Patterns of cortical thickness and surface area in early Parkinson’s disease, NeuroImage 55 (2) (2011) 462–467. doi:10.1016/j.neuroimage.2010.12.043.

[30] A. R. E. Potgieser, A. van der Hoorn, A. M. Meppelink, L. K. Teune, J. Koerts, B. M. de Jong, Anterior temporal atrophy and posterior progression in patients with Parkinson’s disease, Neuro-degenerative diseases 14 (3) (2014) 125–132. doi:10.1159/000363245.

[31] C. Zhou, X.-J. Guan, T. Guo, Q.-L. Zeng, T. Gao, P.-Y. Huang, M. Xuan, Q.-Q. Gu, X.-J. Xu, M.-M. Zhang, Progressive brain atrophy in Parkinson’s disease patients who convert to mild cognitive impairment, CNS neuroscience & therapeutics (2019). doi:10.1111/cns.13188.

[32] H. Braak, K. D. Tredici, U. Rüb, R. A. de Vos, E. N. Jansen Steur, E. Braak, Staging of brain pathology related to sporadic Parkinson’s disease, Neurobiology of Aging 24 (2) (2003) 197–211. doi:10.1016/S0197-4580(02)00065-9.

[33] Y. Y. Fathy, A. J. Jonker, E. Oudejans, F. J. J. de Jong, A.-M. W. van Dam, A. J. M. Rozemuller, W. D. J. van de Berg, Differential insular cortex subregional vulnerability to *α*-synuclein pathology in Parkinson’s disease and dementia with Lewy bodies, Neuropathology and applied neurobiology 45 (3) (2019) 262–277. doi:10.1111/nan.12501.

[34] I. Aracil-Bolaños, F. Sampedro, J. Marín-Lahoz, A. Horta-Barba, S. Martínez-Horta, M. Botí, J. Pérez-Pérez, H. Bejr-Kasem, B. Pascual-Sedano, A. Campolongo, C. Izquierdo, A. Gironell, B. Gómez-Ansón, J. Kulisevsky, J. Pagonabarraga, A divergent breakdown of neurocognitive networks in Parkinson’s Disease mild cognitive impairment, Human brain mapping 40 (11) (2019) 3233–3242. doi:10.1002/hbm.24593.

[35] H. Braak, E. Ghebremedhin, U. Rüb, H. Bratzke, K. Del Tredici, Stages in the development of Parkinson’s disease-related pathology, Cell and Tissue Research 318 (1) (2004) 121–134. doi:10.1007/s00441-004-0956-9.

[36] H. Yang, X. J. Zhou, M.-M. Zhang, X.-N. Zheng, Y.-L. Zhao, J. Wang, Changes in spontaneous brain activity in early Parkinson’s disease, Neuroscience Letters 549 (2013) 24–28. doi:10.1016/j.neulet.2013.05.080.

